# Opposing control of the respiratory brainstem on multiple timescales achieved by transmitter co-release from the locus coeruleus

**DOI:** 10.1101/2025.02.21.639476

**Authors:** Adrienn G. Varga, Brandon T. Reid, Sebastian N. Maletz, Amanda M. Dossat, Erica S. Levitt

**Affiliations:** Department of Pharmacology and Therapeutics, University of Florida College of Medicine, Gainesville FL, USA; Department of Neuroscience, McKnight Brain Institute, University of Florida College of Medicine, Gainesville FL, USA; Breathing Research and Therapeutics Center, University of Florida, Gainesville FL, USA; Edward F. Domino Research Center, Department of Pharmacology, University of Michigan Medical School, Ann Arbor, MI, USA; Department of Anesthesiology, University of Michigan Medical School, Ann Arbor, MI, USA

**Keywords:** Co-transmission, glutamate, noradrenaline/ norepinephrine, Kӧlliker-Fuse nucleus, breathing, pons, arousal circuitry, vesicular glutamate transporter 2

## Abstract

The locus coeruleus (LC) provides widespread noradrenergic (NAergic) modulation throughout the brain to influence a wide range of functions, including breathing. Although both anatomical and physiological evidence supports the involvement of the LC in both the upstream integration and the downstream modulation of breathing, the circuitry behind the latter is unknown. Here, we show that NAergic LC neurons send projections to the Kӧlliker-Fuse nucleus (KF), a critical site in the control of breathing. Long duration activation of NAergic LC neuron terminals in pontine slices induces persistent inhibitory and excitatory NA currents or increases firing rate in postsynaptic KF neurons. Short stimulation on the other hand leads to the VGluT2-dependent release of glutamate that may be co-released with NA in a monosynaptic circuit. Together these results demonstrate that LC neurons can exert flexible, opposing effects on different timescales via glutamatergic and NAergic signaling onto a key respiratory brainstem nucleus.

## Introduction

The co-release of fast acting neurotransmitters and slower-acting neuromodulators from nuclei with a limited number of neurons provides a basis for an increased capacity to differentially and flexibly influence downstream targets on multiple timescales. This has been reported in both dopaminergic^1–4^ and serotonergic^5,6^ circuits in animals ranging from nematodes and flies to rodents. Such neurotransmitter co-release is transforming the general principles of neuroscience and the current theories on how fundamental brain functions operate^7–9^. Yet major questions remain regarding how widespread co-release of transmitters amongst neuromodulatory centers occur and how they might influence functionally diverse downstream targets.

The locus coeruleus (LC) is one of the main sources of noradrenaline (NA) in the brain^10^. The LC contains a small number of densely packed neurons that control behavioral states and integrate an exceptionally wide array of cognitive and homeostatic functions, including breathing^11–14^. Our current understanding of the LC’s enigmatic functional versatility is that in addition to a range of pre- and postsynaptic NAergic receptor subtypes, and a topographical arrangement of functionally diverse neurons with expansive projections to a broad range of target regions, LC neurons also provide multifactorial commands to distinct downstream regions by co-releasing NA with other signaling molecules ^10,11,16,37^. For instance, the LC co-releases NA with dopamine and galanin, all of which bind to G-protein coupled receptors and provide slower-acting postsynaptic modulation^15,17,18^. Recent findings have shown that the LC can also co-release NA with fast-acting neurotransmitters, specifically in an exclusively excitatory LC circuit targeting the parabrachial nucleus^19^. This work demonstrated that in subregions of the parabrachial nucleus, NA and glutamate co-release plays a significant role in directing feeding behaviors through simultaneous activation of α1- and AMPA-receptors, respectively ^19^.

Modern neuroscience techniques have advanced our knowledge regarding how some LC targets are modulated in different conditions and behaviors, yet many fundamental questions remain unanswered. In particular, the LC’s role in the control and integration of breathing rhythms has piqued the attention of many in the field. A relationship between breathing and higher order brain functions has been supported by recent studies, some of them hinting at the LC as one of the critical gateways between the brainstem and cerebral circuits^20–23^. The LC is interconnected with many breathing centers in the brainstem^24–27^, the activity of most LC neurons fluctuates with the respiratory rhythm^28,29^, and >80% of LC neurons are CO_2_ sensitive and contribute to an increased drive to breathe in response to high blood CO_2_ ^30,31^. Additionally, a subset of LC neurons receives projections from the inspiratory rhythm generator, pre-Bӧtzinger complex, that provide a link between breathing and arousal^32^. These data support the LC’s role in relaying breathing-related information to upstream structures, enabling the integration of breathing with higher order processes. Although both anatomical and physiological evidence supports the involvement of the LC in both the upstream integration and the downstream modulation of breathing rhythm, the circuitry behind the latter is unknown. Additionally, whether the integration and control of breathing by the LC is the result of NAergic actions, or the net effect of multiple transmitter systems, is also not known.

Based on the recently discovered ability of LC neurons to co-release NA and glutamate onto the lateral parabrachial nucleus^19^, we tested the hypothesis that LC neurons project to and modulate the nearby key respiratory nucleus, Kӧlliker-Fuse nucleus (KF). The KF is a critical component of the brainstem breathing circuit, that provides essential input to the breathing rhythm- and pattern-generating areas of the medulla and is indispensable to produce ‘normal’ breaths^21^. KF neurons express NAergic receptors, yet the role and origin of NA in the KF is only speculated^33^. Here, we take advantage of retrograde neural tracing, intersectional genetics, *ex vivo* optogenetics and pharmacology to demonstrate that LC neurons send monosynaptic projections to the KF and modulate KF neuron activity both via glutamate and NA release or co-release, acting on both excitatory and inhibitory NA-receptors.

## Results

### Noradrenaline released from the locus coeruleus can excite or inhibit KF neurons

It is known that the LC and KF region are anatomically connected in some mammalian species, but their presence in mice has not been shown to date^25,26^. Additionally, whether these projections form a functional, direct neural circuit, and whether this circuit is excitatory and/or inhibitory, is unknown. To begin to address these questions, we performed retrograde viral tracing to label LC neurons that project to the KF. We unilaterally injected retrograde AAV-hSyn-HI-eGFP-Cre.WPRE.SV40 into the KF of Ai9-tdTomato reporter mice^34^ (**Figure 1A**). The injections resulted in GFP and tdTomato expression in the LC/peri-LC region, as well as sub-coeruleus areas, indicating prominent projections going to the KF from within, and in the proximity of the LC (**Figure 1B**). Next, we used immunohistochemistry to assess the density of projections sent from the specific subpopulation of tyrosine hydroxylase-expressing (TH; NAergic, dopaminergic neuron marker) LC neurons to KF. This approach labeled on average 31±2.62% of NAergic LC neurons (TH+ neurons) with tdTomato, and on average 55±3.29% of all tdTomato labeled neurons were TH+ **(Figure 1C**; n= 4 mice, 5-6 sections/mouse). The labeled neurons were approximately equally dispersed along the rostro-caudal extent of the LC (**Figure 1D**; F(5,16) = 1.15, *p = 0.375* with one-way ANOVA).

**Figure 1.**
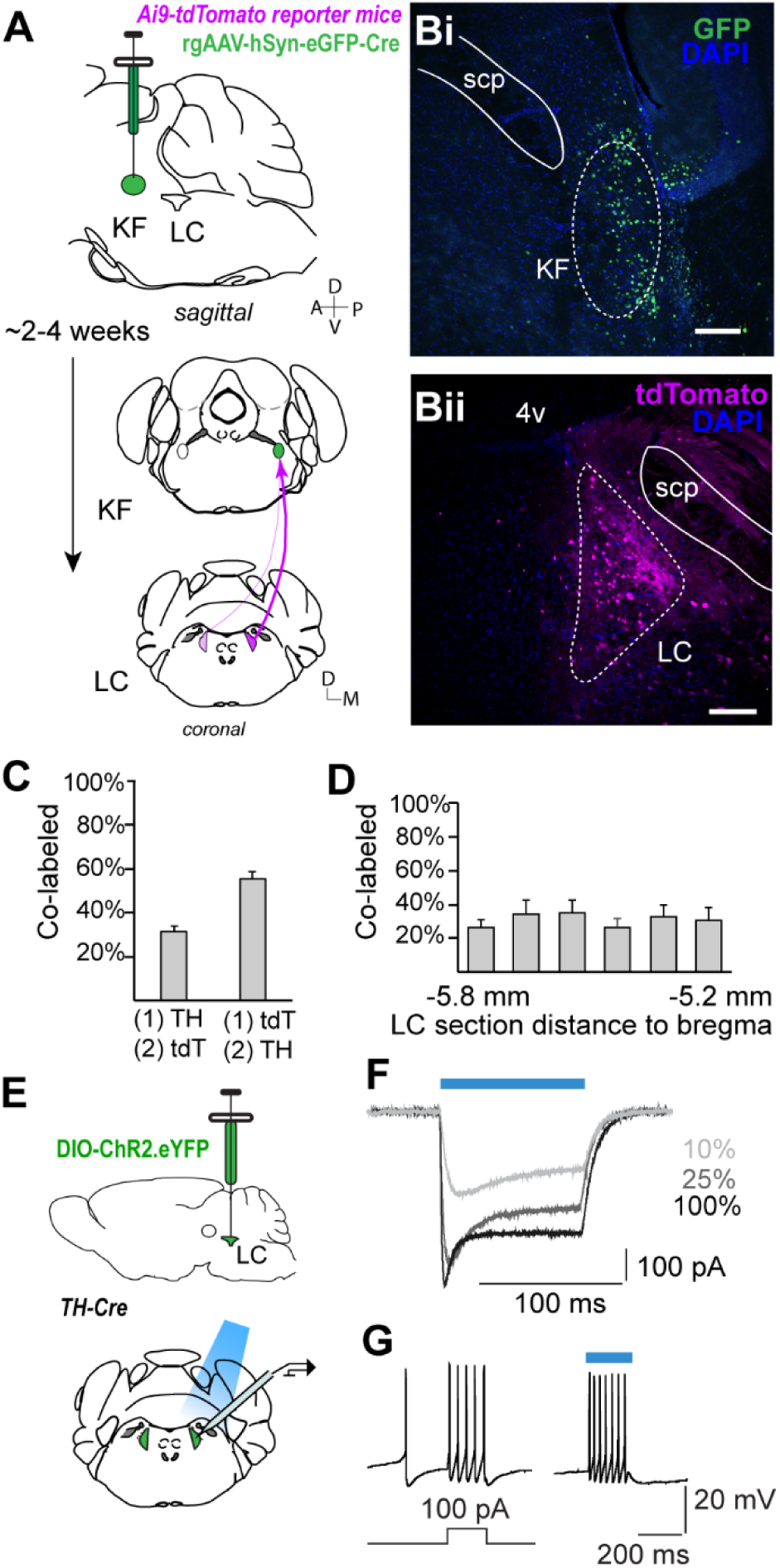
LC neurons send NAergic projections to the breathing center KF. **A** Experimental design for anatomical tracing of LC to KF projections using retrogradeAAV-GFP-Cre injections into the KF of Ai9 tdTomato Cre-reporter mice (*ROSA^LSL-tdT^*). **B** Retrograde Cre-GFP example injection site in the KF (left) and corresponding tdTomato labeled projection neurons in the LC (right). **C** Quantification of co-expression of tdTomato and TH immunofluorescence from Ai9 tdTomato Cre-reporter mice in (A) (n=4, 5-6 sections/mouse). **D** Quantification and localization of tdTomato and TH positive neurons throughout the LC. **E** Experimental design for functional expression of ChR2 in the LC of TH-Cre mice. **F** Voltage clamp recording of an LC NAergic neuron expressing ChR2-eYFP in brain slice, showing inward currents in response to varying intensities of blue light stimulation. **G** Current clamp recordings of action potentials in a ChR2-eYFP expressing neuron following 200ms 100pA current injection and 200 ms blue light stimulation of different intensities (10-100% LED power). Scale bars =100µm.

The LC and the lateral parabrachial nucleus are synaptically coupled in a NAergic-glutamatergic excitatory circuit, that participates in the stress-dependent regulation of feeding^19^. Having established that the KF is also anatomically connected with the LC, we next sought to test whether these areas are also synaptically coupled. We used *ex vivo* optogenetics combined with whole-cell patch clamp recordings in coronal pontine brain slices to begin to elucidate the neurophysiological properties of this circuit. First, we selectively targeted NAergic LC neurons by stereotactically injecting a Cre-dependent AAV to express Channelrhodopsin2-eYFP (AAV5-EF1α-double floxed-hChR2(H134R)-EYFP) bilaterally into the LC of adult TH-Cre mice (**Figure 1E**). We confirmed in a subset of mice that optical stimulation of LC somas in acute coronal brain slices can generate photocurrents and action potentials in response to 470nm light (**Figure 1FG**; n=4 mice, n= 7 slices). Next, we tested whether LC NAergic neurons are functionally coupled with KF neurons, thus we optically stimulated LC axon terminals in KF containing coronal slices while performing whole-cell patch clamp recordings from KF neurons (**Figure 2A**). Given that α2 receptors are present in the KF ^33^, we first sought to test whether NA release induced by light stimulation of LC projections leads to sustained inhibitory currents in KF neurons. NA release from LC to KF was achieved by long duration light stimuli (10sec duration, 20Hz pulse train, or 40Hz pulse train, or continuous light on [data in these groups were statistically similar *p* > 0.05 and were pooled]), and surprisingly, resulted in increased firing rate (**Figure 2BC**; mean normalized firing rate ± SEM: 1191± 887.4 % of baseline; baseline vs. photostimulation; *p = 0.015* with Wilcoxon test), and in many cases (45%, 30 out of 66 neurons, n = 32 mice) in persistent excitatory currents in voltage clamp recordings. These persistent inward currents were blocked by the α1 receptor antagonist prazosin (100nM), confirming α1 receptor expression in the KF for the first time (**Figure 2 DEF**; mean current amplitude ± SEM: baseline 9.56 ± 2.1 pA vs. prazosin 1.96 ± 0.41 pA; *p* = *0.0002* with Mann-Whitney test). These data support the hypothesis of excitatory NAergic modulation of KF neurons by LC.

**Figure 2.**
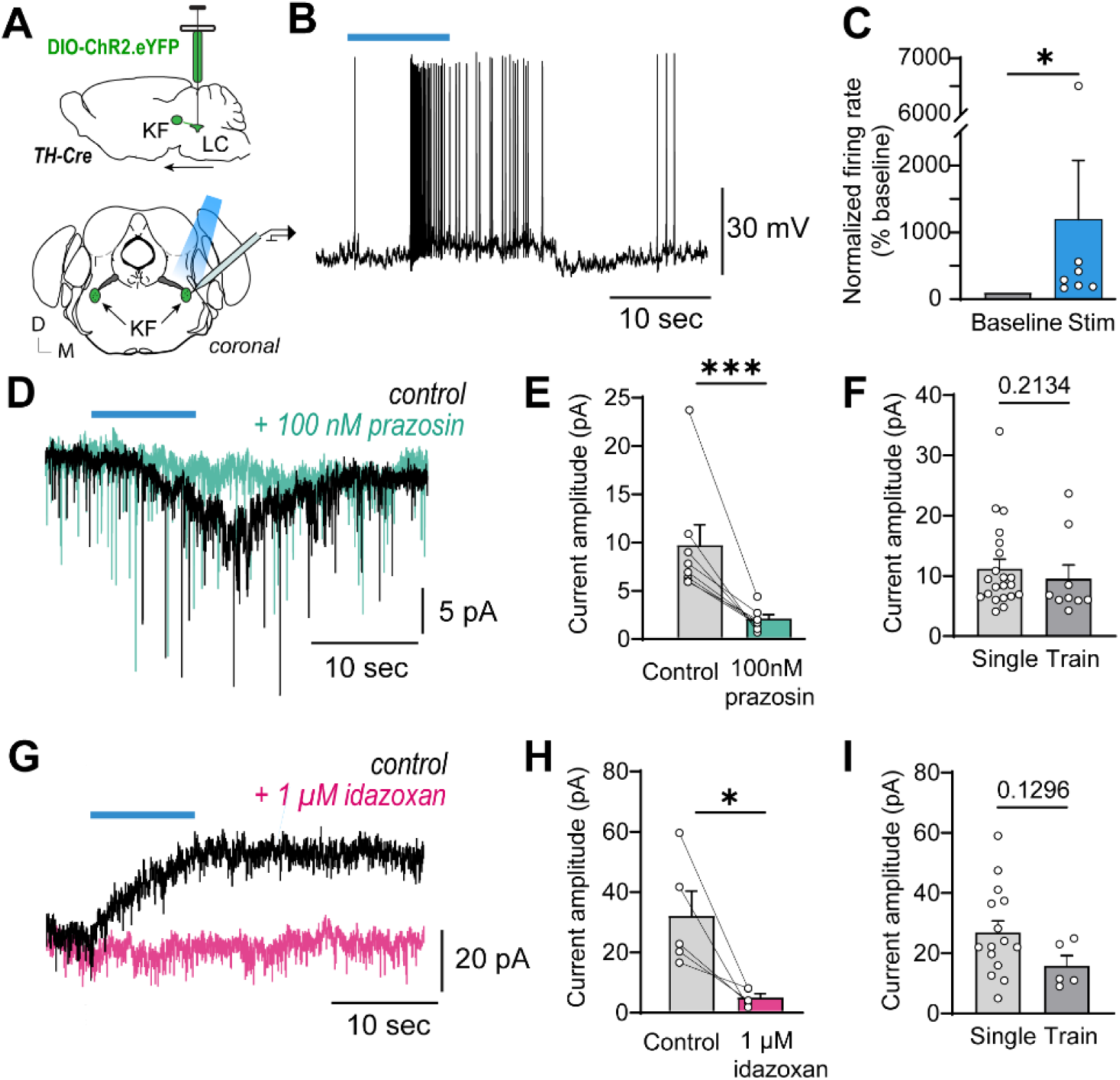
LC neurons can excite or inhibit KF neurons by releasing NA and binding to postsynaptic α1- or α2-receptors. **A** Experimental design for functional mapping of LC to KF projections. ChR2 injected in the LC of TH-Cre mice is efficient to optically excite NAergic LC neuron terminals and induce NA release in pontine KF slices, not containing the LC somas, simultaneous with KF whole-cell recordings. **BC** 10 sec optical stimulation of neurotransmitter release from LC NAergic neurons induces increased KF action potential firing, quantified across cells from n=4 mice. **D-F** Optically evoked persistent inward currents in KF voltage clamp recordings were abolished by the α1-receptor antagonist prazosin (100nM). The amplitudes of the persistent currents were similar regardless of stimulation paradigm (10 sec single continuous light vs. 20 or 40Hz pulse train [pooled]). **G-I** Optically evoked persistent outward currents in KF voltage clamp recordings were abolished by the α2-receptor antagonist idazoxan (1µM). The absolute amplitudes of the persistent currents were similar regardless of stimulation paradigm (10 sec continuous light on vs. 20 or 40Hz pulse train [pooled]). *p<0.05, ***p<0.001 by Wilcoxon test (**C**) or Mann-Whitney (**E, F, H, I**).

As expected, based on anatomical evidence^33^, using the same long duration light stimulus approach as described above, inhibitory currents were induced in 30% of the recordings (20 out of 66 neurons, n = 32 mice). Such α2-receptor mediated persistent inhibitory currents were abolished by the α2 receptor antagonist idazoxan (1 µM) (**Figure 2 GHI;** mean current amplitude ± SEM: baseline 31.59 ± 8.11 pA vs. idazoxan 4.44 ± 1.28 pA; *p = 0.027* with paired two-tailed *t* test). These data provide the first physiological evidence for NAergic signaling in the KF and identifies two subpopulations of KF neurons modulated in opposing manners by the same neuromodulator.

### VGluT2-dependent monosynaptic glutamate release from LC-NA neurons onto KF

LC neurons can co-release both NA and glutamate, but whether this phenomenon only manifests in dual excitatory projections is unknown. While monoamines, such as NA, exert their neuromodulatory effects on longer timescales and frequently via asynaptic mechanisms, glutamate neurotransmission usually requires monosynaptic coupling between the pre- and post-synaptic neurons to achieve rapid signal transmission with both spatial and temporal precision. Therefore, to test the hypothesis that LC neurons are monosynaptically coupled with KF neurons, we used shorter optical stimulation paradigms to elicit glutamate release from LC terminals. First, when stimulated with a 500 ms blue light flash, LC activation resulted in increased KF neuron firing rate in current clamp, that was resistant to the α1 receptor antagonist, prazosin (100nM), demonstrating no or minimal NAergic drive (**Figure 3A**, green trace). However, bath application of the AMPA-receptor antagonist DNQX (100µM) completely blocked action potential firing supporting the role of glutamate in this excitatory pathway (**Figure 3A**, purple trace). When using a short stimulus paradigm (5ms pulse) in voltage clamp, photostimulation led to rapid optically-evoked excitatory postsynaptic currents (oEPSCs; 21 neurons, n =16 mice; **Figure 3B**). The oEPSCs occurred at a short latency following the onset of light flashes (mean latency of oEPSC ± SEM 3.14 ± 0.24 ms) and low jitter (mean SD of latencies ± SEM 0.66 ± 0.09 ms), auspicious of monosynaptic coupling.

**Figure 3.**
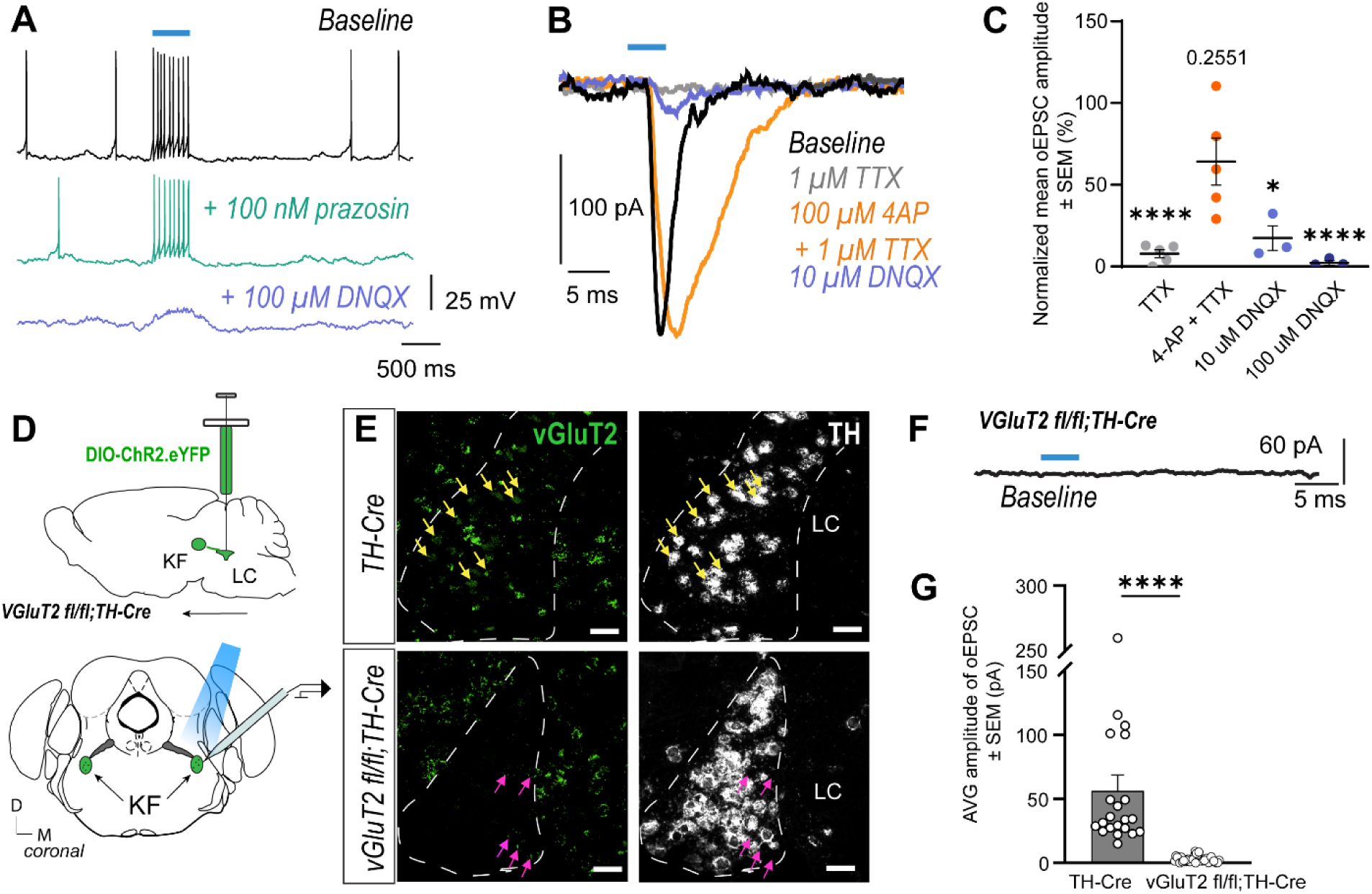
LC neurons directly excite KF neurons via noncanonical glutamate release. **A** Optically evoked neurotransmitter release from LC terminals induces action potential firing in KF neurons that is not sensitive to the α1-receptor antagonist prazosin, but is abolished by the AMPA-receptor antagonist DNQX. **BC** Optically evoked EPSCs are monosynaptic as shown by synaptic isolation of ChR2-dependent events in TTX and 4AP. Monosyanptic oEPSCs are abolished by the AMPA-receptor antagonist DNQX. **D** vGluT2^fl/fl^::TH-Cre mice were injected with ChR2 in the LC to test the contribution of vGluT2 to KF oEPSCs. **E** *vGlut2* mRNA is present in TH-Cre mice, but is eliminated from vGluT2^fl/fl^::TH-Cre mice in the LC region. Yellow arrows: cells co-labeled for vGluT2 and TH mRNA. Magenta arrows: moderate labeling of vGluT2 only. **FG** Optical stimulation of LC terminals in the KF of vGluT2^fl/fl^::TH-Cre mice does not produce any glutamatergic oEPSCs. Scale bars 25µm. *p<0.05, ****p<0.0001 by Mixed effects ANOVA (**C**) or Mann-Whitney test (**G**).

To provide conclusive evidence for monosynaptic coupling, in the next subset of experiments, we bath applied TTX (1µM) to block all Na^+^-dependent neurotransmitter release and isolate ChR2-dependent events which identifies direct excitatory input from LC to KF (**Figure 3BC**). In these recordings oEPSCs were reliably blocked by TTX (mean normalized amplitude ± SEM: 7.83 ± 2.49 % of baseline baseline) and returned close to baseline amplitudes with the co-application of 4 aminopyridine (100µM 4-AP; mean normalized amplitude ± SEM: 64.21 ± 14.35 %), providing pharmacological verification for the monosynaptic junction of LC and KF neurons. Next, to identify the excitatory substance in this synapse, we applied the AMPA-receptor antagonist DNQX (10 or 100 µM), which at the lower 10 µM dose inhibited (mean normalized amplitude ± SEM: 17.38 ± 7.51 %), and at the higher 100 µM dose completely abolished the oEPSCs (mean normalized amplitude ± SEM: 2.18 ± 1.5 %; **Fig 3BC**) confirming glutamate release from LC onto KF.

Although previous work exploring the potential of glutamate release from LC resulted in discordance in the field^35,36^, recently more evidence emerged supporting this phenomenon^19^. Nevertheless, we investigated the origin of the glutamate currents and the possible role of the type 2 vesicular glutamate transporter (vGluT2) in this specific synapse. We generated conditional knockout mice by crossing TH-Cre and vGluT2^flox/flox^ mice to eliminate the machinery necessary for vesicular glutamate release specifically from TH+ neurons in the brain, predicting that ChR2-dependent, light-evoked glutamatergic oEPSCs originating from LC neurons would be absent (**Figure 3DE**). As hypothesized, no glutamatergic events were induced by short light stimulation (5ms pulse) in any of the 39 recorded neurons (**Figure 3FG**; n = 24 mice, mean peak current amplitude ± SEM within 10ms of light onset: TH-Cre mice 56.38 ± 12.21 pA vs. vGluT2^fl/fl^::TH-Cre mice 2.89 ± 0.38 pA; *p < 0.0001* with Mann-Whitney test), bolstering our initial conclusion of the glutamatergic source of the oEPSCs in LC NAergic neurons, and providing further evidence for TH and vGluT2 being co-expressed in a subpopulation of LC neurons, as well as LC neurons relying on vGluT2 for releasing glutamate. Similarly to the TH-Cre mice, vGluT2^fl/fl^::TH-Cre mice showed persistent NAergic currents in the LC to KF circuit (10 / 39 KF neurons, from n = 13 mice), with 40% of the neurons displaying α1-receptor mediated excitatory currents, and the other 60% showing α2-mediated inhibitory events. The amplitudes of the NAergic currents recorded from vGluT2^fl/fl^::TH-Cre mice were statistically similar to the currents recorded from TH-Cre mice (*p* > 0.05 with unpaired *t* test; data were pooled, and are shown in **Figure 2EH**). Together these data support the role of vGluT2 in glutamate release from LC neurons and also demonstrate that this specific transporter does not influence pre-synaptic NA, unlike in another catecholaminergic group of neurons, where vGluT2 facilitates the loading, storage and release of dopamine^39,41^.

### LC neurons co-release NA and glutamate onto the KF

The results described above raise the possibility of NAergic LC neurons to co-release glutamate and NA, and the opportunity for LC neurons to exert both excitatory-excitatory and excitatory-inhibitory postsynaptic modulation. This interplay between the temporally and spatially precise glutamatergic activation and the slowly building, persistent NAergic excitation/inhibition of a downstream target holds great potential for fine-tuning the neural control of widespread LC circuits.

Since dual excitation by glutamate and NA by LC neurons has already been established, we next probed this intricate circuit for coordinated excitatory and inhibitory effects. The experiments depicted in **Figure 5** revealed that short photostimulation (5ms pulse) may lead to glutamatergic oEPSCs and a concomitant persistent outward inhibitory current mediated by inhibitory α2-receptors in the KF (n = 3, from 3 mice; example in **Figure 4A**). Next, bath application of the α2-receptor agonist UK 14304 (1µM) resulted in a persistent outward current that occludes the smaller inhibitory NA currents, if present (**Figure 4AI and II**), meanwhile significantly reducing the amplitude of the glutamatergic oEPSCs (**Figure 4 B**; normalized current amplitude ± SEM: UK14304 27.62 ± 13 % vs. baseline, *F*(1,2) = 20.95, *p = 0.044* by repeated measures one-way ANOVA). Next, bath application of the α2-antagonist idazoxan (1µM), abolished all traces of an α2 outward current and restored the amplitude of oEPSC to baseline (**Figure 4AIII and B**). These data suggest that α2 autoreceptors can inhibit glutamate release from LC terminals.

**Figure 4.**
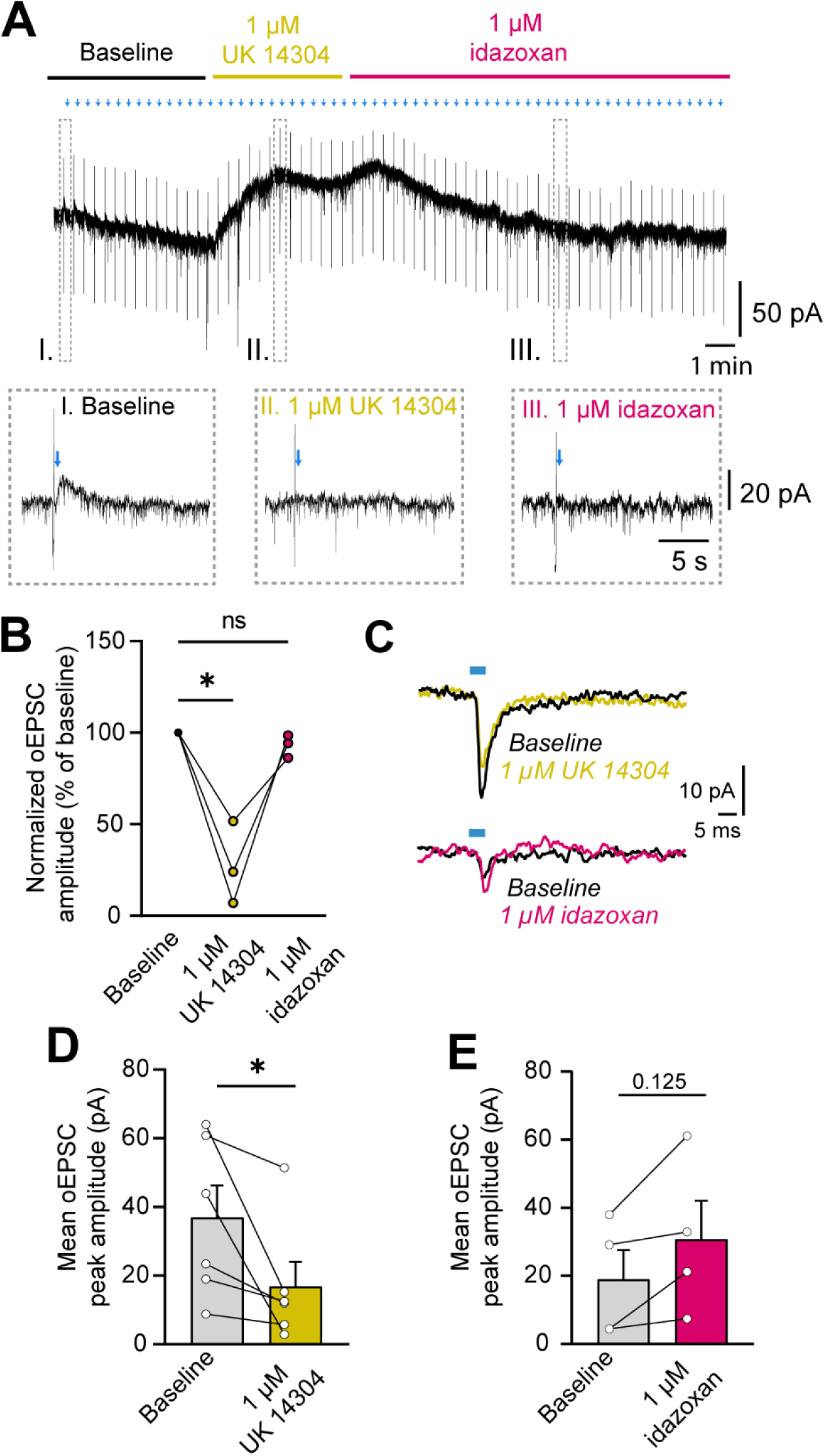
LC neurons co-release NA and glutamate onto KF neurons. **A** Example trace of a KF voltage clamp recording during baseline, α2-receptor agonist UK 14304 (yellow) and α2-receptor antagonist idazoxan (pink) bath application. Blue marks indicate timing of optical stimulation to elicit glutamate release. Boxed traces are zoomed in at the indicated time periods (beginning of baseline[I], near maximal amplitude of α2-current [II] and returning to baseline [III] Note the small outward α2-current in box I. Such evidence of corelease was only present in n=3 recordings, analyzed in the next panel **B** Normalized average glutamatergic oEPSCS amplitude after drug application **C** Example traces of glutamatergic oEPSCs (not from the example recording in A). **D** Bath application of UK 14,304 results in significantly reduced glutamatergic oEPSC amplitude even in recordings where a slow α2-current is not present **E** Bath application of idazoxan without prior UK14304 does not significantly increase oEPSC amplitude. *p<0.05 by RM one-way ANOVA (**B**) or Wilcoxon test **(D,E**).

In a separate set of experiments, where short 5ms optical stimuli only led to glutamatergic, but no apparent NAergic currents, we observed a similar decrease in glutamatergic oEPSC amplitude when the α2-receptor agonist UK 14304 (1µM) was applied (**Figure 4C top trace, D;** mean current amplitude ± SEM: baseline 36.89 ± 9.4 pA vs. UK14304 16.86 ± 7.2 pA, *p = 0.031* by Wilcoxon test).

Taking a similar approach in another cohort, the antagonist idazoxan (1µM) - with no previous UK14304 application - led to a trending increase in glutamatergic oEPSC amplitude (**Figure 4C bottom trace, E;** mean current amplitude ± SEM: baseline 18.99 ± 8.6 pA vs. idazoxan 30.69 ± 11.4 pA, *p = 0.125* with Wilcoxon test), suggesting that endogenous NA tone or co-release of NA minimally affects glutamate release in *ex vivo* brain slices. These results demonstrate the co-release of glutamate and NA from LC to KF, and the multiplexing of opposing signals on multiple timescales exerted from LC-NA neurons onto their target area, in this case, a major breathing center in the brainstem. Incorporating these modalities of LC signaling in future models of neural control by the LC could have a transformative effect on how LC NAergic neurons control a myriad of behaviors and homeostatic function.

## Discussion

The small, tightly packed cluster of LC neurons is responsible for a wide range of cognitive and homeostatic functions^11,37^. More recently, the LC has also become the center of attention in the context of breathing modulation of behavioral states and vice versa, as a potential gateway between respiratory and higher order brain regions. While much is known about the LC’s contributions to the control of breathing^30,31,38,40^, its connections with respiratory nuclei are surprisingly understudied. Yackle et al. (2017) found excitatory input from the pre-Bӧtzinger complex to the LC that is thought to play a role in coordinating arousal states with the breathing rhythm – a potential physiological substrate for the calming effects of breathwork and meditation^32^. On the other hand, the LC can increase respiratory rate through mostly unidentified neural pathways, which likely serve to adjust breathing patterns in state-dependent or voluntary manners^14,31,42^. Here, we describe a novel monosynaptic pathway between LC NAergic neurons and a key respiratory nucleus, the KF. This synapse co-releases glutamate and NA to provide fast glutamatergic excitation and slow NAergic excitation and inhibition of subsets of KF neurons. This circuit is a prime candidate for facilitating the integration of emotional, cognitive, and state-dependent information within the breathing networks of the brainstem to directly drive changes in breathing rate and pattern in a time-efficient, comprehensive and flexible manner.

Recent development of modern neuroscience techniques led to an exponential increase in discoveries focused on the causal role and signaling mechanisms of LC neurons in a variety of contexts^11,37^. It is becoming increasingly evident that the LC’s ability to flexibly ‘multitask’ can be attributed to the multitude of neuromodulators and neuropeptides released and co-released from its core neurons. Additionally, as recently discovered by Yang et al. (2021), NAergic LC neurons also can exert excitatory effects on multiple timescales via NA and glutamate co-release^19^. In the interrogated feeding circuit, they found that LC neurons co-release NA and glutamate to excite parabrachial neurons, which was tested in multiple mouse lines, including conditional VGluT2 knockout mice. The signaling mechanism in this specific LC circuit is directly involved in the fear-induced suppression of feeding, and more importantly, raises the possibility of multi-scale excitation of LC target regions via glutamate co-release. Here, also relying on intersectional genetic methods but utilizing a different pathway and a different conditional knockout mouse line, our findings provide additional physiological evidence for LC neurons co-expressing VGluT2 and TH in adult mice, and again confirm VGluT2’s debated presence, and role in the co-release of glutamate from NAergic LC neurons.

Moreover, to add to the complexity of the LC’s repertoire of postsynaptic commands, our experiments demonstrate NA and glutamate co-release resulting in concomitant inhibition and excitation in different time domains on postsynaptic KF neurons. Interestingly, we did not find dual excitatory effects of co-release in this pathway. Nevertheless, long optical stimulation of transmitter release from LC axon terminals resulted in excitatory α1-receptor mediated persistent currents, without preceding glutamatergic oEPSCs, in a large proportion of the recorded KF neurons. Whether the subpopulation of KF neurons experiencing α1-receptor mediated excitation and the subpopulation mediated by NAergic inhibition and glutamate co-release serve distinct roles in the control of breathing, remains to be tested.

Although the concept of glutamate co-release from NAergic neurons is relatively novel, a wealth of data from other monoamine systems utilizing noncanonical glutamate release may accelerate the elucidation of the functional relevance of this signaling method ^9,43^. For instance, similarly to the observed persistent NA currents, optogenetically evoked dopamine release also affects postsynaptic neurons on a slower and more persistent time scale compared to the rapid transient effects of co-transmitted glutamate^1,3,44^. These temporal differences relate to a precise control exerted by glutamate that may activate single action potentials and convey temporally precise commands, while dopamine influences a delayed, but long-lasting state of increased excitability, allowing other excitatory synapses to induce postsynaptic firing and potentially prompting plasticity^8^. In the current case, glutamate released from LC onto postsynaptic KF neurons could convey precise, time-locked commands, and after a delay, an inhibitory NAergic tone would reduce excitability and the chances of other excitatory synapses influencing KF action potential firing.

If the NAergic system is indeed relying on similar signaling principles as other catecholamine systems, NA and glutamate release may also be subject to frequency-dependent modulation. In the dopaminergic system, in the nucleus accumbens shell, dopamine causes D2-autoreceptor mediated presynaptic inhibition and D1-mediated facilitation of glutamate EPSCs ^44^. At tonic firing frequencies the presynaptic inhibitory effect and the postsynaptic facilitation are counterbalanced, while at burst-firing frequencies postsynaptic facilitation dominates and glutamatergic responses are amplified. Glutamate co-release studies in the dopaminergic system also found that vGluT2 can promote monoamine uptake by the vesicular monoamine transporter 2 (vMAT2) into synaptic vesicles by changing the pH gradient, which provides a mechanism for decreased dopamine release in the absence of vGluT2 ^45^. Our experiments in vGluT2^flox/flox^::TH-Cre mice demonstrated no significant changes in NA current amplitudes, suggesting that glutamate and NA may not be loaded into the same vesicles at this synapse. However, further work determining subcellular mechanisms and machinery of vesicle packaging in NA neurons is needed.

The physiological and functional implications of NAergic signaling in the KF are completely unknown. Glutamatergic signaling on the other hand has been well characterized^33^. Glutamate is the most prevalent neurotransmitter in the KF, participating in the direct modulation of both respiratory rate and pattern^46–49^. AMPA receptors are widely expressed in the KF region, and although NMDA receptors were blocked in our experimental conditions, glutamate released from LC axon terminals is expected to also bind to these receptors. Activation of NMDA receptors in the KF stimulates breathing and influences sleep-wake states, which are also regulated by the LC through largely unexplored mechanisms ^46,50,51^. KF neurons also contain orexin receptors and receive input from the hypothalamus, providing further support for the KF’s role in state-dependent control of breathing^33,52^. Future studies focusing on the intersection of orexin/NA/glutamate modulation in the KF will be an important step in understanding state-dependent and circadian drive in the control of breathing, which is highly relevant for a variety of breathing and sleep disorders.

In summary, our results demonstrate the flexibility, and spatiotemporal diversity of signaling capabilities of LC NAergic neurons, acting in opposing ways via α1 and α2 adrenergic receptors, and tuning NAergic signaling with simultaneous release of glutamate in a subset of LC-KF synapses. While this is a specific pathway with a potentially critical impact on the control of breathing, these findings have global implications for how the field of neuroscience models and evaluates LC “NAergic” signaling in the CNS, whether that is in the context of learning and memory, wakefulness/arousal control, sensory processing, or autonomic regulation by the LC. Additionally, these findings also imply that loss of glutamate release, or disease-induced imbalance of excitatory and inhibitory control of downstream LC targets (including the KF) may contribute to the symptoms and progression of diseases where LC integrity is compromised, such as Alzheimer’s and Parkinson’s disease, or neuropsychiatric disorders.

## Methods

### Animals

Adult mice (male and female, 3-6 months old) were used for all experiments. Mice were bred and maintained in house at the University of Florida. Ai9-tdTomato Cre reporter mice (B6.Cg-Gt(ROSA)26Sor^tm9^(CAG–tdTomato)^Hze^/J; JAX# 007909) ^34^ were used for tracing retrograde tracing experiments. TH-Cre (B6.Cg-7630403G23Rik^Tg^(Th–cre)^1Tmd^/J JAX#: 008601) ^53^ mice were used for brain slice recording and optogenetics experiments and histology. To eliminate the ability of TH+ neurons to release glutamate, TH-Cre mice were crossed with VGluT2^fl/fl^ (Slc17a6^tm1Lowl^/J JAX#: 012898) ^54^ to create VGluT2^fl/fl^::TH-Cre mice, that were used for brain slice recordings with optogenetics and RNAscope. Mice used for brain slice electrophysiology received an intracranial injection between 8-16 weeks of age.

All mice were group-housed in standard sized, individually vented plastic cages (Allentown Jag 75 micro-vent system; Allentown, PA; L: 29.2cm W: 18.5cm H:12.7cm) in pathogen free rooms. Corncob bedding and a NestletTM (Ancare; Bellmore, NY) were in each cage. Mice were housed on a 6:00-18:00hr light cycle with lights on during the daytime, and all experiments were conducted during the light cycle. Temperature in the housing room averaged 70 ± 2°F, with 30-70% humidity and 10-15 room air changes per hour. All animal care was conducted with in the AAALAC animal research program of the University of Florida, in accordance with the guidelines from the Guide for the Care and Use of Laboratory Animals and approved by the University of Florida Institutional Animal Care and Use Committee. We did not see sex differences in our experiments, thus data from both sexes were combined and reported together.

### Stereotaxic injections and viruses

Mice were anesthetized with isoflurane (2-4% in 100% oxygen, Zoetis) and mounted in a stereotaxic frame (Kopf Instruments, Tujunga, CA) where anesthesia was maintained through a nose cone. Body temperature was maintained with a heating pad. Meloxicam (5mg/kg, s.c., Patterson Veterinary) was administered before incision and exposure of the skull. The dorsal skull was then exposed and levelled horizontally between bregma and lambda. A small craniotomy was made to access either the KF (y = -5 mm, x = ±1.7 mm, from bregma) for anatomical tracing experiments, or the LC (y = -5.4 mm, x = ± 0.85 mm) bilaterally for slice electrophysiology. A glass micropipette filled with retrograde AAVrg-hSyn-HI-eGFP-Cre.WPRE.SV40 virus (Addgene, 105540-AAVrg, 7×10¹² vg/mL titer) was lowered into the KF (z = -3.9 mm) of Ai9-tdTomato Cre reporter mice for the retrograde labeling experiments. These mice received 50nl virus unilaterally. For TH-Cre and VGluT2^fl/fl^::TH-Cre mice, the micropipette was filled with AAV5-EF1a-double floxed-hChR2(H134R)-EYFP-WPRE-HGHpA (Addgene, 202298-AAV5, titer 1×10¹³ vg/mL), then lowered into the LC (z = -3.65 mm) and 250 nl of virus was injected bilaterally. Virus was injected with a Nanoject III (Drummond Scientific Company, Broomall, PA, USA) at 50nl/s rate with 20 sec inter-pulse intervals. Following injection, the pipette was left in place for ∼10 min and slowly retracted over the course of at least 5 min. The wound was closed with Vetbond tissue adhesive (3M Animal Care Products, St. Paul, MN, USA). Mice were then placed in a recovery chamber and kept warm until they ambulated normally. Histology experiments with retrograde AAV were performed 4 weeks later, while electrophysiology experiments were performed 6-10 weeks later. Placement of injections was verified in brain slices (50-100 µm or 230 µm) from every mouse by visualizing GFP fluorescence using a multizoom microscope (Nikon AZ100).

### Anatomical tracing and immunohistochemistry

Ai9-tdTomato Cre reporter mice previously injected with retrograde AAV-hSyn-HI-eGFP-Cre.WPRE.SV40 virus in the KF, were deeply anesthetized with isoflurane (Zoetis) 2-4 weeks following surgery, then transcardially perfused with PBS followed by 10% phosphate-buffered formalin. Brains were removed, cryoprotected and stored in 10% formalin/30% sucrose solution at 4°C before further processing. Coronal brain sections were cut at 40 μm. The free-floating sections were stored in Tris-buffered saline (TBS).

Sections containing the LC (5-6 sections/ mouse) were stained for tyrosine hydroxylase to visualize the overlap of TH+ and retrograde GFP/ td-Tomato labeled LC neurons that project to KF. First, sections were rinsed in TBS, then washed in diluting buffer (TBS containing 2% BSA, 0.4% TritonX-100, 1% filtered NGS) for 30 minutes. The sections were blocked in 20% NDS in TBS for 30 minutes to reduce nonspecific staining and immediately incubated in primary antibody for 24 hours (monoclonal anti-tyrosine hydroxylase [mouse; Sigma-Aldrich T1299, lot# 015M4759V], 1:1000 in diluting buffer) at 4°C. Following incubation, sections were washed in diluting buffer and incubated in secondary antibody at room temperature for 2 hours (goat anti-mouse IgG Alexa Fluor Plus 647 in diluting buffer). The sections were washed in TBS, rinsed once in ddH_2_O, then serially mounted with Fluoromount-G DAPI (ThermoFisher) mounting medium. Images were collected on a confocal laser scanning microscope (Nikon A1R) with a 10X objective (N.A. 0.3) and a multizoom microscope (Nikon AZ100) with a 1X objective (N.A. 0.1) and equally processed in Fiji ^55^. Cell counting, to identify NAergic LC neurons that project to the KF, was performed in Image J with the ilastik plugin ^56^.

### Fluorescent *in situ* hybridization (RNAscope®)

Triple fluorescent in situ hybridization using RNAscope® was performed to study the potential co-expression of VGluT2 (SLC17A6) and tyrosine hydroxylase (TH). Mice were deeply anesthetized (100 mg/kg, Fatal Plus, Catalog #078069782, Patterson Veterinary) and transcardially perfused with chilled 0.9% saline. Brains were rapidly removed (<1 min), and flash frozen on 2-methyl butane on dry ice (-20°C). Tissue was encapsulated in Tissue-Plus O.C.T Compound (Catalog #23-730-571, Fisher Scientific), wrapped in foil, and stored at -80°C until further processing. Tissue was equilibrated to -20°C in a cryostat (Leica CM1860) for at least 30 minutes prior to sectioning. Coronal sections were collected at 10 µm thickness on SuperFrost Plus slides (Catalog #12-550-15, Fisher Scientific) and stored at -80°C until further processing. All reagents were purchased from Advanced Cell Diagnostics (ACD, Hayward, CA). Sections underwent fixation, dehydration, and *in situ* hybridization according to the manufacturer’s protocol for Fresh-Frozen tissue. Probes used in this study include: SLC17A6 (Catalog #319171-C1), TH (Catalog #317621-C3). Negative and positive control probes were processed in parallel with the target probes to confirm assay performance and RNA integrity of the samples. Images were collected on a confocal laser scanning microscope (Nikon A1R) with a 10X objective (N.A. 0.3) and equally processed in Fiji ^55^.

### Brain slice electrophysiology

Mice were anesthetized with isoflurane and decapitated. The brain was removed, blocked and mounted in a vibratome chamber (Leica VT 1200S, Leica Biosystems, Buffalo Grove, IL, USA). Coronal slices (230 µm) containing the LC and KF (identified based on anatomical markers and coordinates from Franklin and Paxinos (2008)^57^ were cut in ice-cold artificial cerebrospinal fluid (ACSF) that contained the following (in mM): 126 NaCl, 2.5 KCl, 1.2 MgCl2, 2.4 CaCl_2_, 1.2 NaH_2_PO_4_, 11 D-glucose and 21.4 NaHCO_3_ (equilibrated with 95% O_2_/5% CO_2_). Slices were stored at 32°C in glass vials with equilibrated ACSF. MK801 (10 µM) was added to the cutting solution and for the initial incubation of slices in storage (at least 30 min) to block NMDA receptor-mediated excitotoxicity. Following incubation, the slices were transferred to a recording chamber that was perfused with equilibrated ACSF, warmed to 34°C (Warner Instruments/ Harvard Apparatus, Holliston, MA) and flowing at a rate of 1.5-3 ml/min.

Cells were visualized using an upright microscope (Nikon FN1) equipped with custom built IR-Dodt gradient contrast illumination and DAGE-MTI IR-1000 camera. For recordings performed in the LC, eYFP fluorescence was identified in infected LC cells using LED epifluorescent illumination, eYFP excitation/emission filter cube and detected using a DAGE-MTI IR-1000 camera with sufficient sensitivity in the eYFP emission range.

Whole-cell recordings from were performed with a Multiclamp 700B amplifier (Molecular Devices, Sunnyvale, CA) in voltage-clamp (V_hold_= -60mV) or current clamp mode. Recording pipettes (1.5 – 3 MΩ) were filled with internal solution that contained (in mM): 115 potassium methanesulfonate, 20 NaCl, 1.5 MgCl_2_, 5 HEPES(K), 2 BAPTA, 1-2 Mg-ATP, 0.2 Na-GTP, adjusted to pH 7.35 and 275-285 mOsM. Liquid junction potential (10 mV) was not corrected. Data were filtered at 10 kHz and collected at 20 kHz with pClamp10.7 (Molecular Devices, Sunnyvale, CA), or collected at 400 Hz with PowerLab (Lab Chart version 8; AD Instruments, Colorado Springs, CO). Series resistance was monitored without compensation and remained < 20 MΩ for inclusion. To block inhibitory fast synaptic transmission picrotoxin (100µM) was added to the ACSF. Additional drugs listed in specific experiments were applied by bath perfusion at the indicated concentrations.

Wide-field stimulation of ChR2 with blue light was achieved with an LED (470nm light source, 1.0A, Thorlabs) and directed through an eYFP filter cube and the 40x objective, which produced 8.5 mW/cm^2^ at the slice, confirmed by measurements with a photodiode. To verify efficient functional expression of ChR2 in LC neurons, whole-cell voltage clamp recordings were performed from eYFP-labeled cells in the LC in brainstem slices and stimulated with blue light at increasing intensities while the cell was held at -80 mV (**Figure 1 F**; 10%, 25%, 100% of max intensity, 100ms light pulse). In current clamp, 100pA current injection (200ms) was compared to the action potentials evoked by a 200ms blue light pulse to verify that optical stimulation induces firing in eYFP expressing LC neurons (**Figure 1G**). Control experiments (not shown) were performed on non-eYFP labeled LC neurons in the same slice, which did not display evoked currents or action potentials.

LC axon terminals containing ChR2 were optically stimulated in KF slices during whole-cell recordings of KF neurons (clamped at -80 mV). Pulse trains (10 sec, 20Hz or 40 Hz, 5 ms pulses) or continuous light stimulation (10 sec continuous) were used to achieve presynaptic neuromodulator release and persistent noradrenergic currents in postsynaptic KF neurons. As shown in Figure 2 F and I, there was the optically-evoked persistent currents were statistically similar regardless of whether they were evoked by a pulse train or continuous light, therefore these data were pooled in the graphs in panels E and H. Light-evoked persistent current amplitude was measured in Lab Chart 8 (AD Instruments, Colorado Springs, CO). First, traces were smoothed with a Bartlett window (7 samples) and the peak amplitude was detected as the maximum value in a manually selected window. The maximum value was then subtracted from the mean amplitude of a 30-60 sec baseline immediately preceding the optically-evoked current. For inclusion in further testing and final analysis, baseline persistent currents had to be consistently reproduced with the same stimulus parameters at least twice.

Glutamate release was induced by 5ms light pulses. The glutamate-release stimulating protocol involved 5ms paired light pulses with a 50 ms inter-stimulus interval, however, for the purpose of this study, only the first oEPSC was analyzed, and no paired-pulse effects are noted. Glutamatergic events were analyzed in Clampfit 10.7 (Molecular Devices, Sunnyvale, CA). Each light-evoked oEPSC episode in a sweep (typically 10–50 sweeps per trial or condition) was digitally filtered at 1 kHz with a Gaussian window, then adjusted to baseline immediately before light onset. Single sweep values were averaged together to obtain a mean oEPSC charge for each trial. Average oEPSC amplitudes across conditions were then compared to each other (<2 min after drug administration in the bath).

Average action potential firing rate was calculated using the action potential search mode in Clampfit 10.7 regardless of the stimulus paradigm used. Action potential event frequency during light stimulation was normalized to an equal duration immediately preceding light onset.

### Drugs

All drugs were reconstituted and stored according to the manufacturers’ directions. (+)-MK801 hydrogen maleate (Product # M108), picrotoxin (Product # P1675), DNQX (Product # D0540), 4-aminopyridine (Product # 275875), and idazoxan hydrochloride (Product # 16138) were obtained from Sigma-Aldrich (St. Louis, MO). Prazosin hydrochloride (Cat. # 0623), UK 14,304 tartrate (Cat. No. 2466), and tetrodotoxin (Cat. No. 1069) were purchased from Tocris Biosciences (Minneapolis, MN).

### Statistical Analysis

All statistical analyses were performed in GraphPad Prism version 10. All error bars represent SEM unless otherwise stated. Data with *n* > 8 were tested for normality with Kolmogorov-Smirnov test. Specific statistical tests and *p* values can be found in the figure legends or Results.

## Acknowledgments

This research was supported by Thomas H. Maren Research Excellence Award to A. G. V., NIH grant K99/R00 HL159232 to A. G. V., and NIH grant R01 DA047978 to E. S. L.

## Author contributions

AGV and ESL designed the project, acquired funding and edited the manuscript. AGV performed most experiments and wrote the manuscript. BTR and SNM performed immunohistochemistry, a portion of imaging, and cell counting. AMD performed *in situ* hybridization.

## Resource availability

### Lead contact

Further information and requests for resources and reagents should be directed to and will be fulfilled by the corresponding author, Adrienn G. Varga

### Materials availability

No unique reagents were generated in this study.

### Data and code availability

All data reported in this paper will be shared by the corresponding author upon request. Any additional information required to reanalyze the data reported in this paper is also available upon request.

